# ATAD1 at Mitochondria-ER Contact Sites Facilitates Mislocalized Protein Transfer to the ER

**DOI:** 10.64898/2026.11.22.768386

**Authors:** Shinya Tashiro, Junpei Matsuzaki, Masaya Ono, Tetsuro Yoshimaru, Toyomasa Katagiri, Shunsuke Matsumoto, Toshiya Endo, Yasushi Tamura

## Abstract

Mitochondria-ER contact sites (MERCs) play important roles in cellular functions such as lipid and Ca^2+^ transport and the regulation of mitochondrial dynamics. To better understand the functional diversity of MERCs, it is critical to identify proteins localized at these sites. Here, we employed CsFiND (Complementation assay using fusion of Split-GFP and TurboID) to identify MERC-associated proteins in HeLa cells. Proteomic and cell biological analyses identified ATAD1, a mitochondrial AAA-ATPase, as a MERC-associated protein. ATAD1 facilitates the transfer of GFP-Pex26ΔC, a mislocalized peroxisomal membrane protein, from mitochondria to the ER. GFP-Pex26ΔC is then degraded by the proteasome in a manner dependent on the ER-resident E3 ubiquitin ligase MARCH6. Notably, proteasome inhibition resulted in the accumulation of GFP-Pex26ΔC at MERCs, suggesting that these sites may participate in this quality-control pathway. Together, these findings suggest a potential role for MERCs in the quality control of proteins mislocalized to mitochondria.

## Introduction

Membrane contact sites (MCSs), where organelle membranes are closely apposed, are recognized as platforms for inter-organelle communication. Among the various types of MCSs, mitochondria-ER contact sites (MERCs) are the most extensively studied and have been shown to mediate Ca²⁺ and lipid exchange, coordinate mitochondrial dynamics, and contribute to autophagosome formation and the regulation of apoptosis (1–6). Despite their functional importance, how these diverse processes are regulated at MERCs remains incompletely understood. Moreover, MERCs may harbor additional functions that have yet to be characterized. Defining the molecular composition of MERCs is therefore essential for understanding their full physiological functions.

In mammalian cells, mitochondria-associated ER membranes (MAMs), were first identified in the 1990s through cell fractionation experiments, which revealed that ER subdomains co-precipitated with mitochondria exhibit high phosphatidylserine synthesis activity (7). Subsequently, extensive studies have identified a variety of MERC-forming proteins including Mitofusin 1 and 2 (8, 9), the IP₃R/Grp75/VDAC complex for Ca^2+^ transfer (2), and apoptosis-related complexes involving Bap31, Fis1, and Tom40 (6, 10). VAP family proteins residing in the ER also contribute to MCS formation by binding FFAT- or FFNT-motif-containing proteins on opposing organelles (11–14). In yeast, ER–mitochondria contact sites are primarily mediated by the ER–mitochondria encounter structure (ERMES) complex, which consists of four core subunits, Mmm1, Mdm10, Mdm12, and Mdm34 (15) Structural and biochemical studies have demonstrated that ERMES mediates phospholipid transfer between membranes (15–19). More recently, in situ cryo-electron tomography revealed that ERMES forms bridge-like structures spanning the ER and mitochondrial membranes, providing structural insights into the mechanism of lipid transfer at MERCs (20). PDZD8, a mammalian SMP domain-containing protein functionally related to the yeast ERMES component Mmm1, has been shown to interact with FKBP8 on mitochondria and Rab7 on late endosomal/lysosomal membranes (21–24). Furthermore, PDZD8 interacts with Protrudin at ER–endolysosome contact sites and exhibits lipid extraction activity, suggesting its involvement in lipid transport and endosome maturation (25).

The MERC proteome has been investigated through large-scale approaches, including proteomic profiling of purified mitochondria-associated membrane (MAM) fractions (26, 27) and proximity labeling-based proteomic approaches (28–32). These studies have identified numerous MERC-associated proteins, including SYNJ2BP and its interacting partner RRBP1 (29), RTN1A (28), and FUNDC2 and MTFR1 (30), highlighting the molecular diversity of MERCs and their potential roles in various cellular processes.

To identify proteins associated with MERCs, we applied CsFiND (Complementation Assay Using Fusion of Split-GFP and TurboID), a proximity-labeling method we previously developed in yeast, to human cultured cells. CsFiND combines split-GFP-based visualization of MCSs with split-TurboID-mediated biotinylation of proteins in their vicinity (33, 34). A key advantage of CsFiND is that proximity labeling occurs specifically at physiologically formed MERCs, whose formation can be simultaneously confirmed by GFP fluorescence, thereby minimizing artifacts associated with artificial tethering (33). Using CsFiND in HeLa cells, we identified a broad set of candidate MERC-associated proteins, including previously known MERC components, supporting the specificity and utility of CsFiND for profiling proteins associated with MERCs in human cells. Among the proteins identified, we focused on ATAD1, an AAA-ATPase localized to the mitochondrial outer membrane (MOM) and the mammalian ortholog of yeast Msp1(35, 36). In yeast, Msp1 extracts mislocalized tail-anchored proteins from the MOM and facilitates their transfer to the ER via the GET pathway, where they are ubiquitinated by the ER-resident E3 ligase Doa10 and subsequently degraded by the proteasome (37, 38). Notably, tMsp1 and Pex15Δ30, a model substrate mislocalized to the MOM, were shown to form complexes predominantly at MERCs, suggesting that substrate transfer to the ER may occur at these sites (37). However, the functional involvement of MERCs in this pathway and whether a similar mechanism operates in mammalian cells remain unclear.

In this study, we show that ATAD1 is associated with MERCs in HeLa cells and facilitates the transfer of GFP-Pex26ΔC, a model protein mislocalized to the MOM, from mitochondria to the ER. Our analyses revealed that GFP-Pex26ΔC is transferred to the ER in a manner partially dependent on ATAD1 and subsequently degraded by the proteasome through a pathway involving the ER-resident E3 ubiquitin ligase MARCH6. Despite its role in mitochondrial-to-ER transfer, ATAD1 was not essential for the overall degradation of GFP-Pex26ΔC, suggesting the involvement of additional pathways. Upon proteasome inhibition, GFP-Pex26ΔC accumulated at MERCs, suggesting that these contact sites may participate in the transfer and quality control of mislocalized mitochondrial proteins. Furthermore, depletion of selected MERC-associated proteins altered the steady-state levels of GFP-Pex26ΔC, supporting a potential functional connection between MERCs and this quality-control process. Together, these findings suggest a previously unrecognized role for MERCs in mitochondrial protein quality control, although whether these contact sites are required for protein transfer to the ER remains to be established.

## Results

### Selection of CsFiND proteins exhibiting the highest biotinylation activity in HeLa cells

CsFiND is a complementation system based on the fusion of split-GFP and split-TurboID fragments, enabling simultaneous visualization of membrane contact sites (MCSs) and proximity-dependent biotinylation of MCS-associated proteins (Fig. 1A) (33). In our previous work using the CsFiND system in yeast, we observed that the efficiency of CsFiND fragment reconstitution varied depending on the combination of split-GFP and split-TurboID fragments (33, 34). To determine the optimal combination, we generated eight different CsFiND construct pairs and assessed their biotinylation activity in HeLa cells. The constructs contained the N-terminal regions of Tom70 and ERj1, including their respective transmembrane domains, which we previously demonstrated to be effective in targeting split-GFP fragments to the MOM and ER, respectively (39).

**Figure 1.**
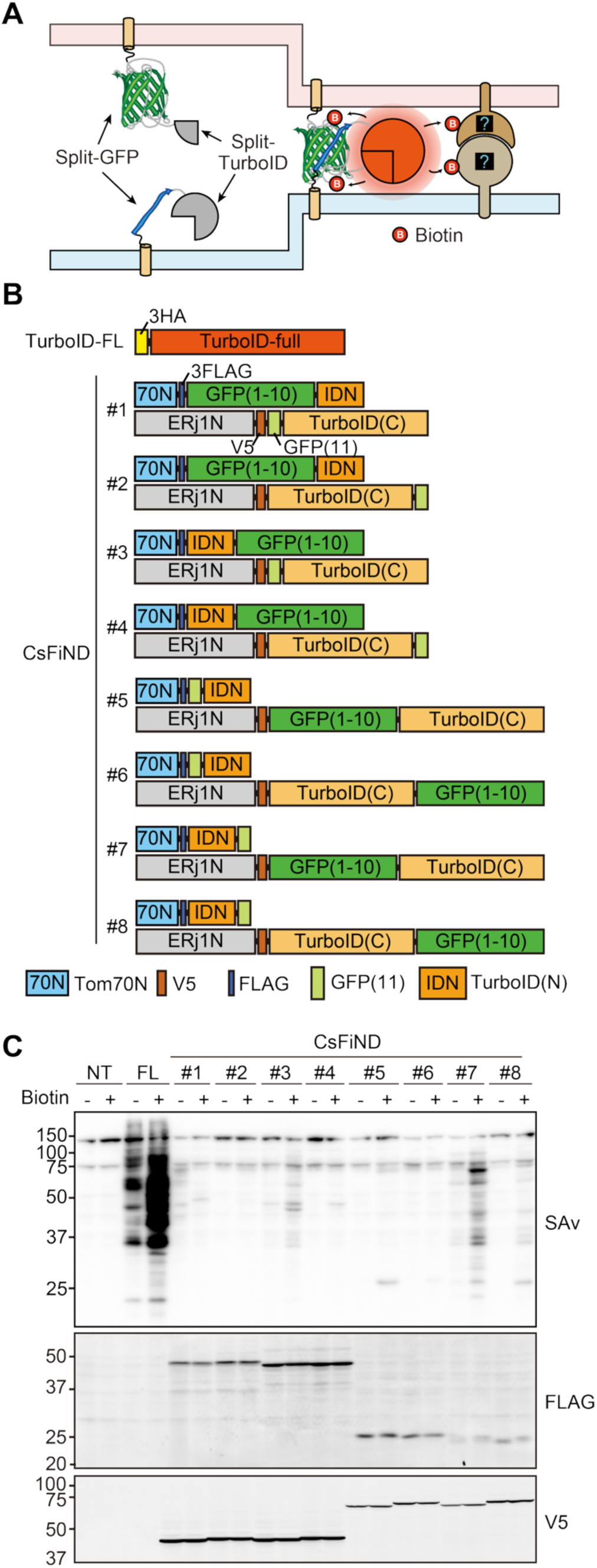
CsFiND for identification of MERC-localized proteins. (A, B) Schematics of the CsFiND experiment labeling MERC-localized proteins and the CsFiND protein pairs used in this study. (C) HeLa cells transiently expressing the indicated CsFiND proteins were incubated for 1 h with or without 50 µM biotin, and total lysates were analyzed by streptavidin-Cy5 (SAv) for biotinylated proteins and by anti-FLAG and anti-V5 antibodies for CsFiND proteins. NT, no transfection; FL, full-length TurboID.

The eight pairs differed in the combinations and arrangements of the split-GFP and split-TurboID fragments (Fig. 1B). HeLa cells were transiently transfected with each of the eight CsFiND construct pairs or with full-length TurboID, and then cultured in biotin-supplemented medium. Immunoblotting of total cell lysates with anti-V5 or anti-FLAG antibodies detected bands corresponding to the expected molecular weights of the CsFiND constructs, confirming their expression (Fig. 1C). Although the expression levels of all CsFiND constructs were generally comparable, the extent of protein biotinylation varied among the tested pairs. Cells expressing full-length TurboID exhibited strong biotinylation signals, whereas those expressing CsFiND proteins showed substantially lower levels of protein biotinylation. Among the eight CsFiND construct pairs tested, pair #7 exhibited the highest biotinylation activity and was therefore used in subsequent experiments.

### Optimization of CsFiND Conditions to Prevent Excessive Membrane Contact Formation

To determine whether CsFiND construct #7, which exhibited the highest biotinylation efficiency, was reconstituted specifically at mitochondria-ER contact sites (MERCs), we monitored the reconstituted GFP signal. Under the same conditions used for biotinylation assays (Fig, 1C), in which equal amounts of plasmids encoding the mitochondria- and ER-targeted CsFiND proteins were co-transfected, the GFP signal appeared to uniformly cover the entire surface of the mitochondria. This result suggested that irreversible reconstitution of split-GFP under these conditions induced excessive formation of MERCs. In our previous study using split-GFP probes to visualize MERCs in HeLa cells, we found that reducing the expression level of the mitochondria-targeted split-GFP suppressed excessive contact formation (39, 40). Based on this observation, we decreased the amount of plasmid encoding the mitochondrial CsFiND protein while keeping the amount of the ER-targeted plasmid constant. As the relative dosage of the mitochondrial construct was reduced, the intensity of the GFP signal decreased and shifted from a widespread pattern along the mitochondria to discrete punctate signals confined to subregions of the mitochondria (Fig. 2A). To assess biotinylation activity under more physiologically relevant conditions, we transfected HeLa cells with the mitochondrial CsFiND construct at a ratio of 1:30 or 1:50 relative to the ER-targeted construct and cultured them in biotin-containing medium. After a 1-hour incubation, cells expressing full-length TurboID exhibited robust biotinylation, whereas those expressing CsFiND proteins showed only faint biotinylation signals, consistent with the limited amount of reconstituted TurboID at MERCs. However, after 24 hours of biotin labeling, biotinylated proteins became detectable in cells expressing CsFiND proteins. Although the signal intensity remained approximately one-tenth of that observed after 1-hour labeling in full-length TurboID-expressing cells, the prolonged labeling appeared to provide sufficient biotinylation for proteomic identification of candidate MERC-associated proteins.

**Figure 2.**
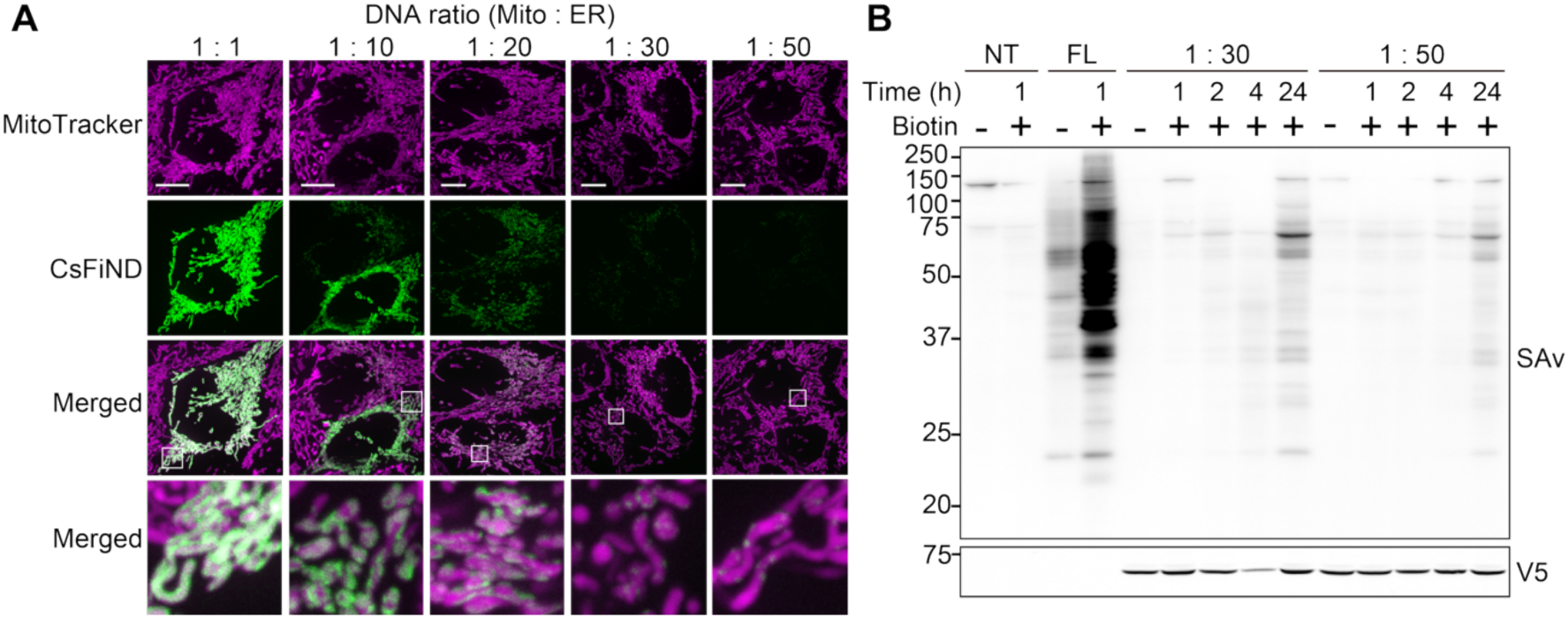
Optimization of CsFiND conditions for proximity labeling. (A) HeLa cells were transfected with plasmids encoding the ER- and mitochondria-targeted CsFiND proteins at ratios ranging from 1:1 to 1:50. Cells were stained with MitoTracker and observed by fluorescence microscopy. (B) HeLa cells that were either non-transfected, transiently expressing full-length TurboID, or transfected with CsFiND plasmids at 1:30 or 1:50 ratios were incubated with or without biotin for the indicated times. Total lysates prepared from these cells were analyzed with streptavidin-Cy5 (SAv) or anti-V5 antibodies.

### ATAD1 is localized at ER–Mitochondria Contact Sites

To identify proteins associated with MERCs, we employed the CsFiND system using CsFiND #7 constructs at the optimized expression ratio (1:30). Total cell lysates were prepared from HeLa cells expressing CsFiND fusion proteins, full-length TurboID, or left untransfected, followed by streptavidin-based purification. To compensate for the lower biotin labeling efficiency of CsFiND, we used ten times more CsFiND-expressing and non-transfected cells than TurboID-expressing cells for purification of biotinylated proteins. Immunoblot analysis of the purified fractions revealed only trace levels of biotinylated proteins in non-transfected cells. In contrast, biotinylated proteins were detected in both the CsFiND- and TurboID-expressing cells, with distinct biotinylation patterns, consistent with spatially restricted biotinylation by CsFiND at MERCs (Fig. 3A). The purified biotinylated proteins were analyzed by LC-MS/MS to identify proteins associated with MERCs. We selected candidate MERC-associated proteins based on at least twofold enrichment in CsFiND samples relative to both negative controls (untransfected cells and cells expressing cytosolic TurboID), with p < 0.05, and visualized the results using volcano plots (Fig. 3B and S1). This analysis identified 207 and 195 significantly enriched proteins relative to non-transfected and TurboID-expressing cells, respectively, with 124 proteins overlapping between the two datasets (Fig. 3C). Notably, the majority of these 124 overlapping proteins were annotated as either ER- or mitochondria-localized, and are highlighted in blue and green, respectively (Fig. 3C). Proteins previously identified as MERC constituents in large-scale proteomic studies are shown in bold (Fig. 3C). Additionally, proteins boxed in yellow represent those that were previously reported to localize at MERCs based on low-throughput studies (Fig. 3C). In the volcano plots, the previously reported MERC proteins marked by magenta dots clustered in the upper-right region, further supporting the specificity of CsFiND for profiling MERC-associated proteins.

**Figure 3.**
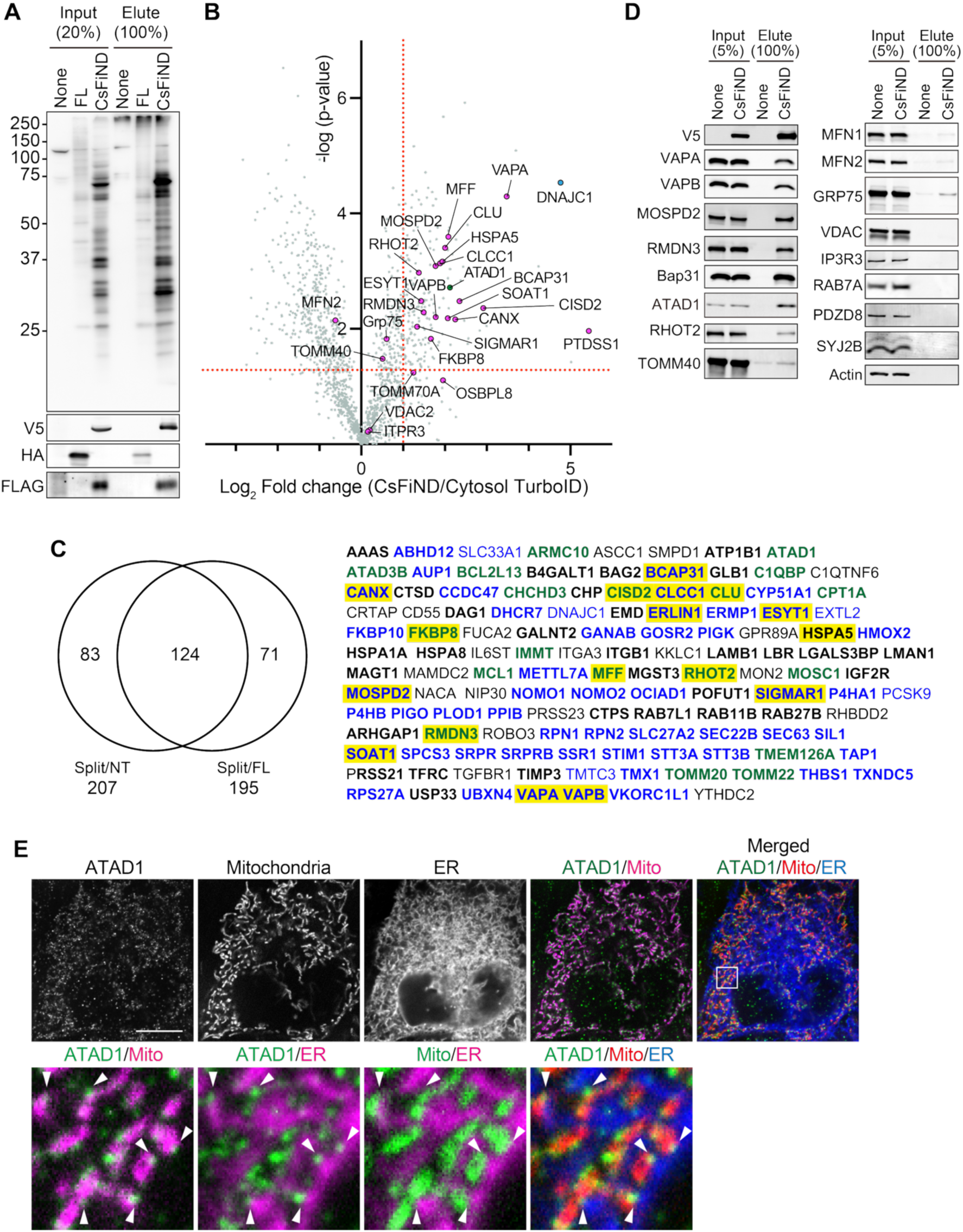
Identification of MERC-associted proteins using CsFiND. (A) Biotinylated proteins were purified from total lysates of HeLa cells using streptavidin beads. None, non-transfected HeLa cells; FL, HeLa cells expressing full-length TurboID; CsFiND, HeLa cells transfected with CsFiND plasmids (1:30). Cells were incubated with biotin for 1 h in the None and FL conditions, and for 24 h in the CsFiND condition. (B) A volcano plot shows the relative abundance of proteins biotinylated in CsFiND-expressing cells compared with full-length TurboID-expressing cells. Factors previously reported to localize at MERCs in individual studies (not large-scale proteomic analyses) are highlighted in magenta. (C) The number of biotinylated proteins enriched (Log2 fold change ≥ 1, p < 0.05) in CsFiND-expressing cells, compared with either non-transfected cells (Split/NT) or full-length TurboID-expressing cells (Split/FL), is shown. A total of 124 proteins were common to both comparisons and are listed on the right. ER and mitochondrial proteins are shown in blue and green, respectively. MERC proteins identified in large-scale analyses or individual studies are shown in bold and boxed in yellow, respectively. (D) Biotinylated proteins from HeLa cells with or without expression of CsFiND proteins were purified with streptavidin and analyzed by immunoblotting using the indicated antibodies. (E) Immunofluorescence analysis of ATAD1 localization. HeLa cells expressing Su9-BFP as a mitochondrial marker and mCherry-Sec61b as an ER marker were stained with an anti-ATAD1 antibody.

Although DNAJC1 showed the highest level of enrichment, this likely reflects its use as a component of the CsFiND fusion construct. PTDSS1, the first protein reported to be enriched at MERCs, was also recovered (41). Besides, several well-established MERC proteins were robustly and reproducibly enriched in CsFiND samples. These included VAPA, VAPB, and MOSPD2, which recognize FFAT motifs through their MSP domains. Another identified protein was RMDN3, a MOM protein that interacts with VAPB via an FFAT-like motif (11–14). Immunoblotting also confirmed the biotinylation of these factors, further supporting the reliability of CsFiND for identifying MERC-associated proteins (Fig. 3D). Taken together, these findings demonstrate that CsFiND enables spatially resolved profiling of proteins associated with MERCs. Taken together, these findings demonstrate that CsFiND enables spatially resolved profiling of proteins associated with MERCs.

Among the enriched proteins, we focused on ATAD1, the mammalian ortholog of yeast Msp1 (36, 42). In our previous study, we used split-GFP to visualize the proximity between Msp1 and Pex15Δ 30, a model tail-anchored protein mislocalized to mitochondria, and found that the resulting GFP puncta were frequently observed at MERCs in yeast (37). To test whether ATAD1 localizes to MERCs, we examined the subcellular localization of ATAD1 by immunofluorescence microscopy (Fig. 3E). ATAD1 was detected as discrete puncta distributed along the tubular mitochondria labeled by Su9-BFP. Some of these puncta were located adjacent to the ER, visualized using mCherry-Sec61β, suggesting that ATAD1 partially localizes to MERCs.

### ATAD1 facilitates the transfer of mislocalized mitochondrial proteins to the ER

To investigate whether ATAD1 is involved in the clearance of proteins mislocalized to mitochondria, we first established a model substrate in mammalian cells. In yeast, the peroxisomal tail-anchored membrane protein Pex15 is known to mislocalize to mitochondria when it lacks its C-terminal 30 amino acids (Pex15ΔC30) (37). We therefore examined the subcellular localization of Pex26, the mammalian ortholog of yeast Pex15, in HeLa cells. We transiently expressed GFP-tagged full-length Pex26 or a truncated version lacking the C-terminal 30 amino acids (GFP-Pex26ΔC) and analyzed their localization by fluorescence microscopy (Fig. 4A). GFP-Pex26 exhibited punctate signals that colocalized with the peroxisomal marker Pex3-mCherry, confirming its localization to peroxisomes (Fig. 4B). In contrast, GFP-Pex26ΔC colocalized with the mitochondrial marker Tom20, indicating that deletion of the C-terminal region causes Pex26 to mislocalize to mitochondria (Fig. 4B).

**Figure 4.**
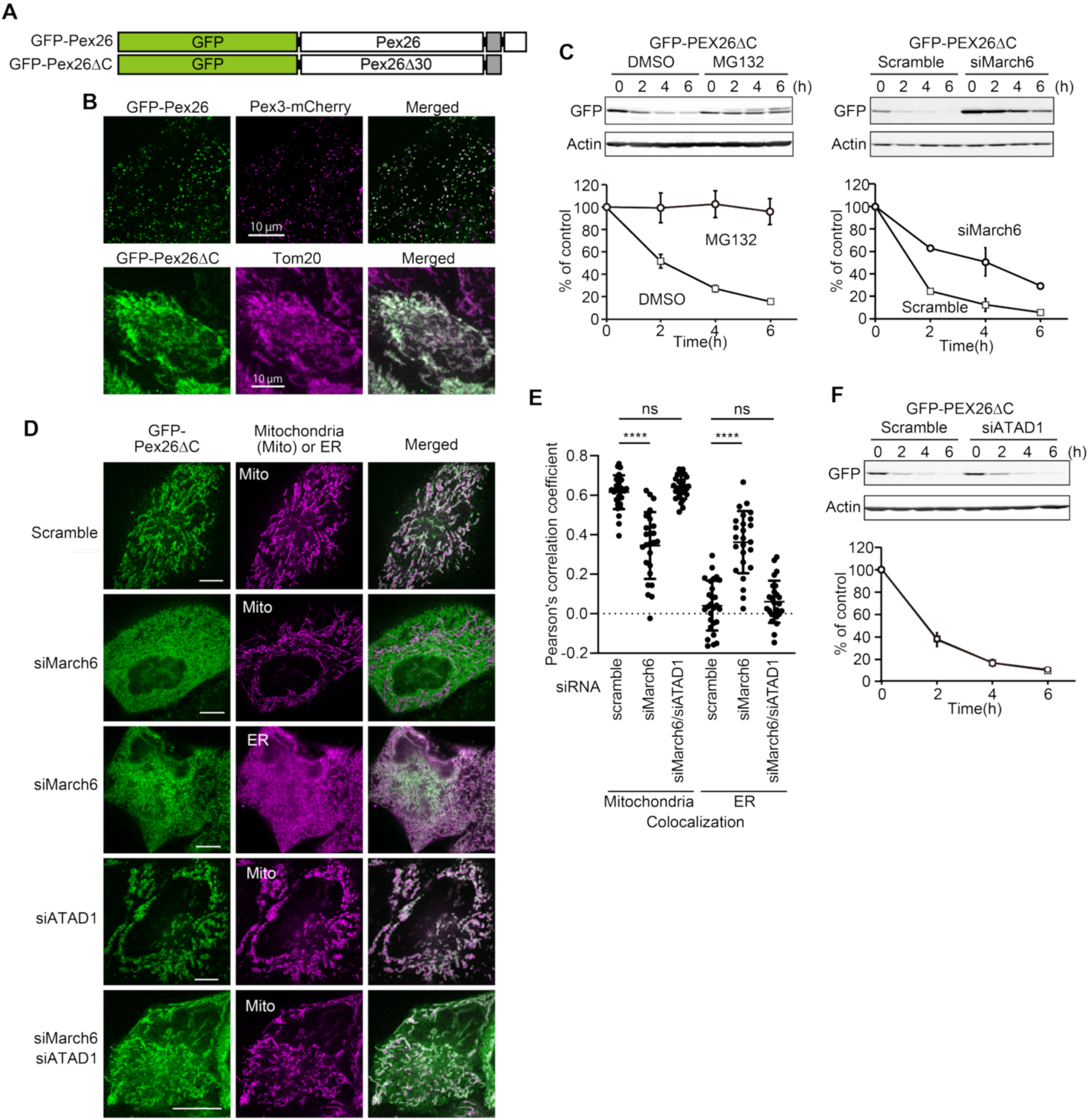
GFP-Pex26ΔC is degraded by the proteasome in a MARCH6-dependent manner. (A) Model substrate protein mislocalized to mitochondria. (B) GFP-Pex26 and Pex3-mCherry were transiently expressed and visualized by fluorescence microscopy. GFP-Pex26ΔC-expressing cells were analyzed by immunofluorescence using an anti-Tom20 antibody. (C) HeLa cells stably expressing GFP-Pex26ΔC were treated with 50 μg/mL cycloheximide and incubated for the indicated times in the presence of DMSO, 10 µM MG132, or scramble siRNA or MARCH6 siRNA. Total cell lysates were analyzed by immunoblotting with anti-GFP and anti-Actin antibodies. The line graph shows the degradation kinetics of GFP-Pex26ΔC. The amount of GFP-Pex26ΔC at time 0 after cycloheximide treatment was set to 100%. Data are presented as mean ± SEM. (n = 3). (D) HeLa cells stably expressing GFP-Pex26ΔC were treated with the indicated siRNAs and analyzed by confocal microscopy. Mitochondria and the ER were visualized with MitoTracker and mCherry-Sec63, respectively. (E) Pearson’s correlation of GFP-Pex26ΔC colocalization with mitochondria or the ER. Lines indicate mean ± SD. (n = 25). Statistical significance was assessed using one-way ANOVA followed by Dunnett’s multiple comparisons test. ns, not significant, ***p < 0.001. (F) HeLa cells stably expressing GFP-Pex26ΔC were treated with 50 μg/mL cycloheximide and incubated for the indicated times in the presence of scramble siRNA or ATAD1 siRNA. Total cell lysates were analyzed by immunoblotting with anti-GFP and anti-actin antibodies.

We next examined whether GFP-Pex26ΔC is recognized as a mislocalized protein and degraded by the proteasome, as reported for Pex15ΔC30 in yeast (37). HeLa cells expressing GFP-Pex26ΔC were treated with the proteasome inhibitor MG132, and protein stability was assessed by cycloheximide chase experiments. MG132 treatment markedly stabilized GFP-Pex26ΔC, indicating that GFP-Pex26ΔC is degraded in a proteasome-dependent manner (Fig. 4C). To identify an E3 ubiquitin ligase involved in this process, we focused on MARCH6, the mammalian ortholog of the yeast E3 ligase Doa10, which ubiquitinates Pex15ΔC30 (37). siRNA-mediated knockdown (KD) of MARCH6 significantly stabilized GFP-Pex26ΔC and delayed its degradation, indicating that MARCH6 contributes to the degradation of GFP-Pex26ΔC (Fig. 4C).

To determine whether GFP-Pex26ΔC is transferred from mitochondria to the ER before degradation, we examined its subcellular localization following MARCH6 KD. Whereas GFP-Pex26ΔC was predominantly localized to mitochondria in cells treated with scramble siRNA, MARCH6 KD resulted in its accumulation at the ER, as indicated by increased colocalization with the ER marker mCherry-Sec63 (Fig. 4D, E). These results suggest that GFP-Pex26ΔC is transferred from mitochondria to the ER, where MARCH6 contributes to its proteasomal degradation, and that impairment of this degradation pathway leads to the accumulation of GFP-Pex26ΔC at the ER.. We next asked whether ATAD1 contributes to the mitochondrial-to-ER transfer of GFP-Pex26ΔC. In MARCH6-KD cells, GFP-Pex26ΔC predominantly accumulated at the ER, whereas simultaneous KD of MARCH6 and ATAD1 shifted its localization toward mitochondria (Fig. 4D, E). Similar results were obtained in ATAD1-Knock out (KO) cells (Fig. S2). However, a fraction of GFP-Pex26ΔC remained associated with the ER even in the absence of ATAD1, indicating that ATAD1 facilitates, but is not essential for, the transfer of GFP-Pex26ΔC from mitochondria to the ER (Fig. 4D, S2).

We further examined whether ATAD1 depletion affects the overall degradation of GFP-Pex26ΔC. Despite its effect on the subcellular localization of GFP-Pex26ΔC, ATAD1 depletion did not delay its degradation (Fig. 4F). Thus, although ATAD1 contributes to the mitochondrial-to-ER transfer of GFP-Pex26ΔC, it does not appear to be essential for its overall degradation. Together with the residual ER localization of GFP-Pex26ΔC observed in the absence of ATAD1, these results suggest that ATAD1-independent mechanisms also contribute to the transfer and/or subsequent degradation of GFP-Pex26ΔC.

### GFP-Pex26ΔC accumulates at mitochondria–ER contact sites upon proteasome inhibition

Given the localization of ATAD1 at MERCs, we investigated whether GFP-Pex26ΔC is transferred from mitochondria to the ER via MERCs. To this end, we examined the subcellular localization of GFP-Pex26ΔC following inhibition of proteasomal degradation with MG132. Under these conditions, GFP-Pex26ΔC exhibited a distinct punctate distribution, in contrast to its predominant mitochondrial localization under normal conditions and its ER localization in MARCH6-KD cells (Fig. 5A). Strikingly, these puncta were associated with both the ER and mitochondria, suggesting that stabilized GFP-Pex26ΔC accumulates at MERCs. To further examine the localization of GFP-Pex26ΔC at MERCs, we employed hypotonic swelling to enlarge the ER and mitochondrial membranes and facilitate visualization of their contact sites (43). Under these conditions, GFP-Pex26ΔC accumulated at membrane contact interfaces between ER and mitochondrial vesicles (Fig. 5B).

**Figure 5.**
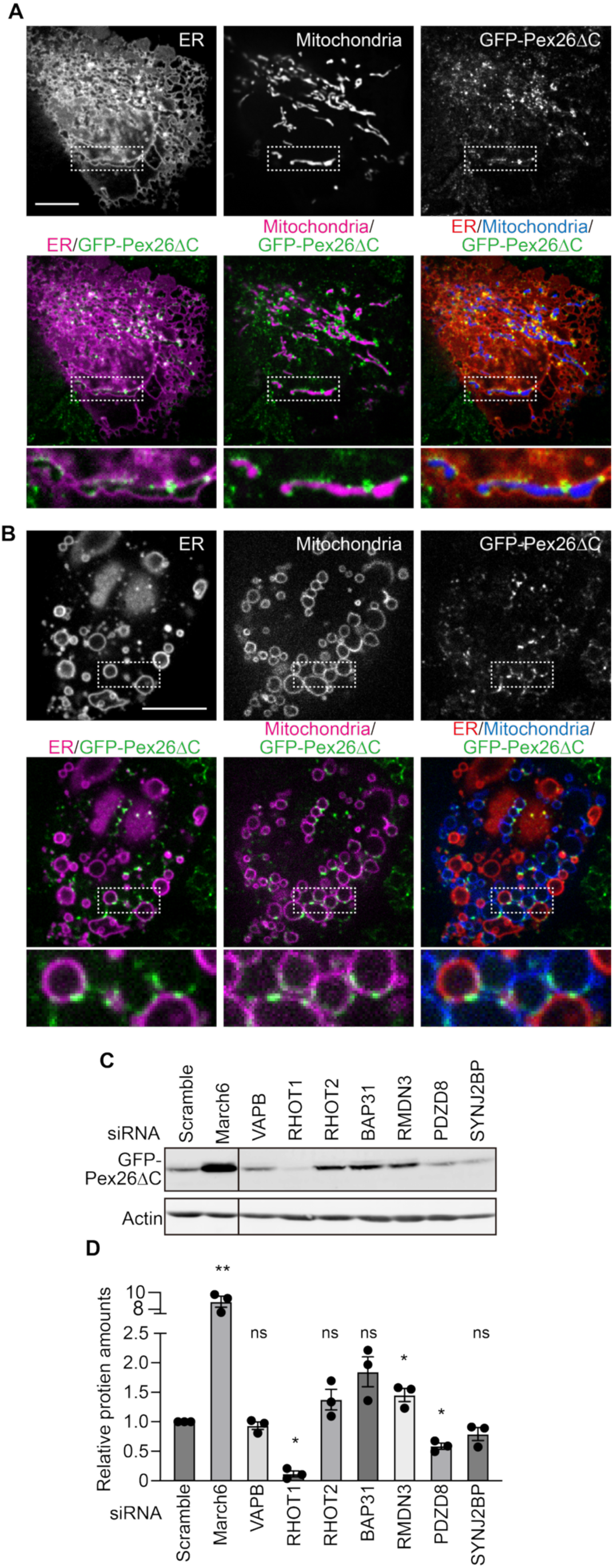
GFP-Pex26ΔC accumulates at MERCs upon proteasome inhibition. (A) HeLa cells stably expressing GFP-Pex26ΔC were transfected with plasmids expressing mCherry-Sec63 (ER marker) and Su9-BFP (mitochondrial marker), treated with 10 µM MG132 for 8 h, and analyzed by confocal microscopy. (B) The same cells as in (A) were subjected to hypo-osmotic treatment to vesiculate the ER and mitochondria. (C) Total cell lysates were prepared from cells treated with the indicated siRNAs and analyzed by immunoblotting with anti-GFP and anti-actin antibodies. (D) GFP-Pex26ΔC levels were quantified and normalized to actin, with the level in cells treated with scramble siRNA set to 1. Bar graphs represent the mean ± SEM from three independent experiments (n = 3). Statistical significance was assessed using two-tailed one-sample t-tests on log2-transformed normalized values against a hypothetical mean of zero. ns, not significant, *p < 0.05; **p < 0.01.

We next asked whether MERCs are functionally involved in the clearance of GFP-Pex26ΔC. To this end, we individually depleted MARCH6 and several proteins previously reported to localize to MERCs, including VAPB (11), RHOT1 (44), RHOT2 (45), BAP31 (6), RMDN3 (11), PDZD8 (21), and SYNJ2BP (29), and examined the steady-state levels of GFP-Pex26ΔC by immunoblotting of total cell lysates. MARCH6 KD resulted in a marked accumulation of GFP-Pex26ΔC, whereas none of the other factors tested had a comparable effect (Fig. 5C and D). Among the MERC-associated proteins, RMDN3 KD significantly increased GFP-Pex26ΔC levels, while BAP31 KD showed a similar trend without reaching statistical significance. In contrast, RHOT1 KD reduced GFP-Pex26ΔC levels, whereas PDZD8 knockdown caused a smaller decrease. KD of VAPB, RHOT2, or SYNJ2BP had little or no detectable effect. These results suggest that several MERC-associated proteins may influence the steady-state levels of GFP-Pex26ΔC, although none had an effect comparable to that of MARCH6.

## Discussion

In this study, we applied the CsFiND system to human cells for spatially resolved proteomic profiling of MERCs. CsFiND recovered a number of well-established MERC proteins, including VAPA, VAPB, MOSPD2, and RMDN3, supporting the specificity and utility of the method. However, several known MERC factors, such as Mfn2, PDZD8, and SYNJ2BP, were not enriched. SYNJ2BP overexpression nevertheless increased MERC formation in our experiments, consistent with a previous report (29). These observations suggest that some MERC proteins may be inefficiently detected by CsFiND because of limited accessibility to the CsFiND labeling enzyme. MERC composition may also vary depending on cell type or physiological conditions. Application of CsFiND to different cell types and under different physiological conditions should therefore provide a more comprehensive view of MERC composition and dynamics.

Among the proteins identified by CsFiND, we focused on ATAD1, a mitochondrial AAA-ATPase and the mammalian ortholog of yeast Msp1. Our results suggest that a subset of ATAD1 localizes to MERCs (Fig. 3E) and that ATAD1 facilitates the transfer of GFP-Pex26ΔC, a model protein mislocalized to mitochondria, from mitochondria to the ER (Fig. 4). GFP-Pex26ΔC was degraded in a proteasome-dependent manner, and depletion of the ER-resident E3 ubiquitin ligase MARCH6 delayed its degradation and caused its accumulation at the ER (Fig. 4C, D). Notably, simultaneous depletion of ATAD1 and MARCH6 shifted GFP-Pex26ΔC localization from the ER toward mitochondria, supporting a role for ATAD1 in facilitating its transfer to the ER. However, ATAD1 depletion did not significantly affect the overall degradation kinetics of GFP-Pex26ΔC (Fig. 4F). This finding suggests that ATAD1-dependent transfer to the ER is not essential for the degradation of this substrate. One possibility is that ATAD1-independent mechanisms also mediate the transfer of GFP-Pex26ΔC to the ER, allowing its subsequent degradation. Alternatively, GFP-Pex26ΔC may be degraded through additional pathways that do not require its transfer to the ER. These possibilities are not mutually exclusive, and the relative contributions of different pathways remain to be determined.

How ATAD1 promotes the transfer of extracted proteins to the ER remains unclear. In yeast, the GET system has been shown to function downstream of Msp1-mediated extraction and to facilitate the transfer of extracted membrane proteins to the ER (38). Whether the mammalian GET pathway plays a similar role downstream of ATAD1 remains unknown. Identifying the machinery that receives ATAD1-extracted proteins and transfers or inserts them into the ER membrane will therefore be important for understanding the mechanism of mitochondrial-to-ER protein transfer at MERCs.

A central question is how ATAD1 recognizes proteins that should be removed from mitochondria. Rather than recognizing specific sequence motifs, ATAD1 may preferentially act on orphan membrane proteins that remain unassembled or lack stable binding partners. In this model, proteins that fail to form protective complexes would be more accessible to ATAD1, regardless of whether they are bona fide mitochondrial proteins or proteins mislocalized from other organelles such as the Golgi apparatus (36). Native mitochondrial proteins may be relatively resistant to ATAD1 because their assembly into protein complexes protects them from extraction. Such a mechanism would allow ATAD1 to remove mislocalized or unassembled proteins without requiring strict substrate specificity.

Our findings also raise the question of whether MERCs are functionally required for the transfer and degradation of mislocalized mitochondrial proteins. Proteasome inhibition caused GFP-Pex26ΔC to accumulate at sites where mitochondrial and ER membranes were closely apposed (Fig. 5A, B), suggesting that MERCs may participate in this process. One possibility is that MERCs facilitate the transfer of proteins extracted from mitochondria to the ER by bringing the two membranes into close proximity. However, the accumulation of GFP-Pex26ΔC at MERCs does not necessarily demonstrate that its transfer occurs through these sites or that MERCs are essential for this process. For example, GFP-Pex26ΔC may accumulate at MERCs because its degradation is inhibited, without these sites being required for its transfer to the ER. Furthermore, the persistence of GFP-Pex26ΔC degradation in ATAD1-deficient cells raises the possibility that alternative pathways can compensate for the loss of ATAD1-dependent transfer. Directly examining whether disruption of MERCs impairs the mitochondrial-to-ER transfer of GFP-Pex26ΔC will therefore be important for establishing the functional contribution of these contact sites.

Our analysis of MERC-associated proteins further suggests that several factors may influence GFP-Pex26ΔC turnover. Depletion of RMDN3 modestly but significantly increased the steady-state level of GFP-Pex26ΔC, whereas BAP31 depletion showed a similar trend that did not reach statistical significance. In contrast, RHOT1 depletion reduced GFP-Pex26ΔC levels, as did PDZD8 depletion to a lesser extent, whereas depletion of VAPB, RHOT2, or SYNJ2BP had little or no detectable effect. These differential effects suggest that MERC-associated proteins may influence GFP-Pex26ΔC turnover through distinct mechanisms. However, changes in steady-state protein levels do not necessarily reflect changes in degradation rates and could also result from altered protein synthesis, localization, or other cellular processes. Moreover, the effects of individual MERC proteins on GFP-Pex26ΔC levels do not establish whether their functions depend on MERC formation. Further studies examining how these factors affect MERC organization, mitochondrial-to-ER protein transfer, and substrate degradation will be necessary to clarify their roles in this pathway.

Another unresolved question is how ATAD1 localizes to MERCs. ATAD1 forms puncta along mitochondria, a subset of which is juxtaposed with the ER (Fig. 3E). ER-localized interacting proteins or specific membrane environments at MERCs may promote its accumulation at these sites. Identifying the factors responsible for ATAD1 localization to MERCs may help clarify how ATAD1-mediated extraction is coupled to subsequent transfer to the ER.

Taken together, our findings identify ATAD1 as a MERC-associated protein that facilitates the transfer of a mislocalized mitochondrial membrane protein to the ER, where MARCH6 contributes to its proteasomal degradation. The accumulation of GFP-Pex26ΔC at MERCs upon proteasome inhibition further suggests that these contact sites may participate in this quality-control process. However, ATAD1 is not essential for the overall degradation of GFP-Pex26ΔC, and whether MERCs are required for its transfer to the ER remains to be established. Our study therefore suggests a potential role for MERCs in the quality control of mislocalized mitochondrial proteins and provides a basis for investigating the mechanisms that coordinate protein extraction, inter-organelle transfer, and degradation.

## Materials and Methods

### Cell Culture

HeLa cells were cultured in high-glucose Dulbecco’s modified Eagle’s medium (DMEM, FUJIFILM Wako Pure Chemical Corporation, Cat. No. 044-29765) supplemented with 10% fetal bovine serum (FBS) (Nichirei Biosciences Inc., Cat. No. 173012; Lot No. BCBX4446) and 1% penicillin–streptomycin (FUJIFILM Wako Pure Chemical Corporation, Cat. no. 168-23191) at 37°C in a humidified atmosphere containing 5% CO2.

### Plasmids

A synthetic gene encoding 3×HA-TurboID codon-optimized for expression in mammalian cells was synthesized by Eurofins Genomics and cloned into the BamHI/XbaI sites of pcDNA3.1 to generate pMM475.

To generate CsFiND expression constructs, pMM82 (pcDNA3.1-Tom70N-GFP1-10) (40) expressing the N-terminal 70 amino acids of human Tom70 fused to 3×FLAG and GFP1–10, and pMM86 (pcDNA3.1-ERj1N(1–200)-V5-GFP1–10) (39) expressing the N-terminal 200 amino acids of human ERj1 fused to V5 and GFP1–10 were used as backbone vectors. These plasmids were digested with NotI and XbaI to remove the GFP1–10 coding sequence, which was replaced with DNA fragments encoding split-GFP (GFP1–10 or GFP11) and split-TurboID (sTurboIDn or sTurboIDc) fused in different combinations and arrangements using the In-Fusion cloning system. The split-GFP fragments were amplified by PCR from pMM87 (pcDNA3.1-ERj1N-V5-GFP11) and pMM82 (pcDNA3.1-Tom70N-FLAG-GFP1–10), whereas the split-TurboID fragments were amplified from pMM475.

To generate plasmids for lentiviral expression of GFP-Pex26 and GFP-Pex26ΔC, DNA fragments encoding full-length human Pex26 and its C-terminally truncated variant lacking the last 30 amino acids (Pex26ΔC) were amplified by PCR using HeLa cell-derived cDNA as a template. The PCR products were cloned into the XhoI/ApaI sites of the lentiviral expression vector pLVSIN-AcGFP1-C1 (Takara Bio) to generate pMM505 (pLVSIN-AcGFP1-hPex26) and pMM506 (pLVSIN-AcGFP1-hPex26ΔC), respectively, which encode N-terminally AcGFP1-tagged Pex26 and Pex26ΔC.

To generate an ATAD1-targeting sgRNA expression plasmid, lentiCRISPRv2-Blast (a gift from Brett Stringer; Addgene plasmid #98293; RRID:Addgene_98293) (46) was digested with BsmBI and ligated with annealed oligonucleotides containing the ATAD1-targeting sequence (forward, 5′-CACCGACACTGACCGATAAGTGGTA-3′; reverse, 5′-AAACTACCACTTATCGGTCAGTGTC-3′), resulting in pKO95. All plasmids used in this study are listed in Table S1.

### Lentivirus production and transduction

Lentiviral particles were produced in Lenti-X 293T cells (Takara Bio, Cat. no. 632180) by co-transfection of the lentiviral transfer vector with the packaging plasmid psPAX2 (a gift from Didier Trono; Addgene plasmid #12260) and the envelope plasmid pMD2.G (a gift from Didier Trono; Addgene plasmid #12259) using PEI MAX (Cosmo Bio Co., Ltd., Cat. No. 24765-100). For each 35-mm dish, 3 μg of total plasmid DNA and 9 μg of PEI MAX were separately diluted in 150 μL of Opti-MEM (Thermo Fisher Scientific, Cat. no. 31985062). The two solutions were mixed, incubated for 30 min, and added to the cells. At 48 h post-transfection, culture supernatants containing lentiviral particles were collected and filtered to remove cellular debris. HeLa cells were transduced with the resulting lentiviral supernatants in the presence of polybrene and selected with 2 μg/mL puromycin.

### Purification of biotinylated proteins and LC-MS/MS analysis

For proximity biotin labeling, biotin was added to the culture medium at a final concentration of 50 µM. Cells expressing CsFiND constructs were labeled for 24 h, whereas cells expressing full-length TurboID were labeled for 1 h. Cells were lysed in 1 mL of RIPA buffer, and the lysates were clarified by centrifugation at 15,000 × g for 10 min at 4°C. Biotinylated proteins were enriched by incubation of the clarified lysates with Pierce Streptavidin Magnetic Beads (Thermo Fisher Scientific, Cat. no. 88816) for 24 h at 4°C. The beads were washed twice with RIPA buffer, once with 1 M KCl, once with 0.1 M Na2CO3, once with 2 M urea in 10 mM Tris-HCl (pH 8.0), and twice again with RIPA buffer, as described previously (30).

Bead-bound proteins were digested with trypsin, and the resulting peptides were purified using a phase-transfer method and C18 StageTips. Peptides were analyzed using a nanoLC-Ultra 2D system coupled to a TripleTOF 5600 mass spectrometer (AB SCIEX), equipped with a C18 reversed-phase column (75 µm × 150 mm; ChromXP C18-CL, Eksigent). LC-MS data were processed using 2DICAL for peak detection, alignment, and quantification, as described previously (47). Peptides were identified using MASCOT (version 2.5.1; Matrix Science) against the UniProt/Swiss-Prot human protein database. Protein-level abundance values were calculated by averaging the 2DICAL intensities of peptides assigned to each protein. Statistical analyses were performed using R (version 2.7.0).

### siRNA transfection

HeLa cells cultured in 24-well plates (IWAKI, Cat. No. 3820-024) were transfected with siRNA at a final concentration of 30 nM using Lipofectamine RNAiMAX (Thermo Fisher Scientific, Cat. No. 13778030). For each well, 15 pmol of siRNA and 1 μL of Lipofectamine RNAiMAX were separately diluted in 50 μL of Opti-MEM, combined, and incubated for 15 min at room temperature before addition to the cells. Cells were analyzed 72 h after transfection.The siRNAs targeting MARCH6, ATAD1, VAPB, BAP31, PDZD8, and SYNJ2BP were used for gene knockdown. The sequences of all siRNAs are listed in Table S3.

### Immunoblotting

HeLa cells cultured in 48-well plates were directly lysed in 60 μL of SDS sample buffer after washing with 200 μL of PBS. Proteins were separated by SDS-PAGE and transferred onto polyvinylidene fluoride (PVDF) membranes (Immobilon-FL; Millipore, Cat. No. IPFL00010). Membranes were probed with the indicated primary antibodies, followed by Cy5-conjugated goat anti-mouse IgG (Jackson ImmunoResearch Laboratories, RRID: AB_2338714) or DyLight 800-conjugated goat anti-rabbit IgG (Rockland Immunochemicals, RRID: AB_11183542). All primary and secondary antibodies used in this study are listed in Table S2. Fluorescence signals were detected and analyzed using a Typhoon imager (GE Healthcare).

### Immunofluorescence microscopy

HeLa cells grown in 35 mm glass-bottom dish (Matsunami Glass Ind., Ltd., Cat. No. D141410) were fixed for 12 min at room temperature with pre-warmed 4% PFA in phosphate buffer (Nacalai Tesque, Inc. Cat. no. 09154-56) and washed three times with PBS(-) (FUJIFILM Wako Pure Chemical Corporation, Cat. No. 045-29795). Cells were permeabilized with 0.5% Triton X100 in PBS for 10 minutes and washed three times with PBS. After blocking with 3% BSA containing PBS for 1 hour, cells were incubated with Rabbit polyclonal antibody against ATAD1 (Proteintech, 16882-1-AP), Mouse monoclonal antibody against TOM20 (BD transduction laboratories, Cat. No. 612278), in blocking buffer for 1 hour. Cells were washed three times with PBS and incubated with 2 μg/ml goat anti-mouse IgG (H + L) AlexaFluor405 Secondary Antibody, (Thermo Fisher Scientific, RRID: AB_221604) in blocking buffer for 1 hour. Cells were observed under Olympus IX83 microscope with a CSU-X1 confocal unit (Yokogawa), a 100×, 1.4 NA, objective (UPlanSApo, Olympus) and an EM-CCD camera (Evolve 512; Photometrics) manipulated by Metamorph software (Molecular Devices).

### Live cell fluorescence microscopy

HeLa cells grown in 35-mm glass-bottom dishes were transfected with plasmids encoding the CsFiND system as described above. For mitochondria staining, cells were incubated with 100 ng/mL MitoTracker Red CMXRos (Thermo Fisher Scientific, Cat.no. M46752) in Opti-MEM for 5 min at 37°C in 5% CO2. Cells were washed twice with DMEM supplemented with 10% FBS and subjected to microscopic observation. Cells were observed using an Olympus IX83 microscope equipped with a CSU-X1 confocal unit (Yokogawa), a 100×/1.4 NA objective (UPlanSApo, Olympus), and an EM-CCD camera (Evolve 512; Photometrics), controlled using MetaMorph software (Molecular Devices).

### Hypotonic Cell Treatment

To induce morphological changes in mitochondria and the endoplasmic reticulum (ER), HeLa cells were subjected to hypo-osmotic shock by incubation for 10 min in DMEM diluted to 5% (v/v) with water and supplemented with 10 mM HEPES. Cells were subsequently subjected to microscopic observation.

## ACKNOWLEDGMENTS

We are grateful to the members of the Tamura laboratory for helpful discussion. This work was supported by JSPS KAKENHI (Grant Number, 19H03174, 23K23832, and 25K02220 to YT, 20H04912, 20H05689, 20H05929, 25H00978, and 25K21752 to TE), AMED-PRIME (Grant Number JP20gm5910026) from Japan Agency for Medical Research and Development, AMED to YT, and AMED-CREST (Grant Number 21gm1410002), and Takeda Science Foundation grant to TE and YT. ST was a JSPS fellow.

## AUTHOR CONTRIBUTIONS

Conceptualization, S.M., T.E., and Y.T; Methodology, S.T., J.M M.O. and Y.T.; Investigation, S.T., J.M., M.O., and Y.T.; LC-MSMS sample preparation, T.Y., and T.K. Writing – Original Draft, Y.T.; Writing – Review & Editing, S.T., J.M., S.M., T.E., and Y.T.; Funding Acquisition, S.T., T.E., and Y.T.

---

**Fig. S1.**
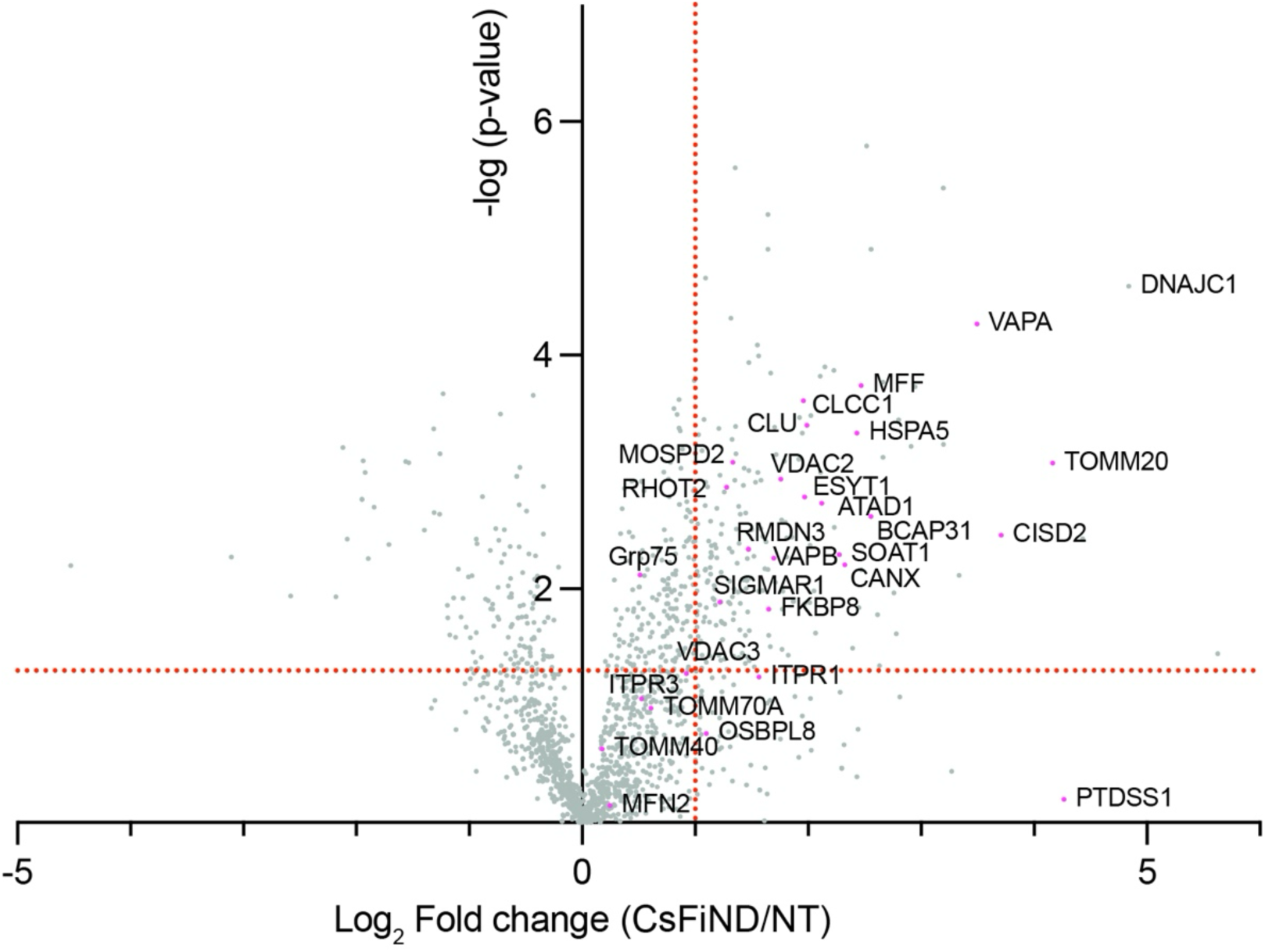
Proteomic profiling of MERC-associated proteins using CsFiND. A volcano plot shows the relative abundance of proteins biotinylated in CsFiND-expressing cells compared with non-transfected cells. Factors previously reported to localize at MERCs are highlighted in magenta.

**Fig. S2.**
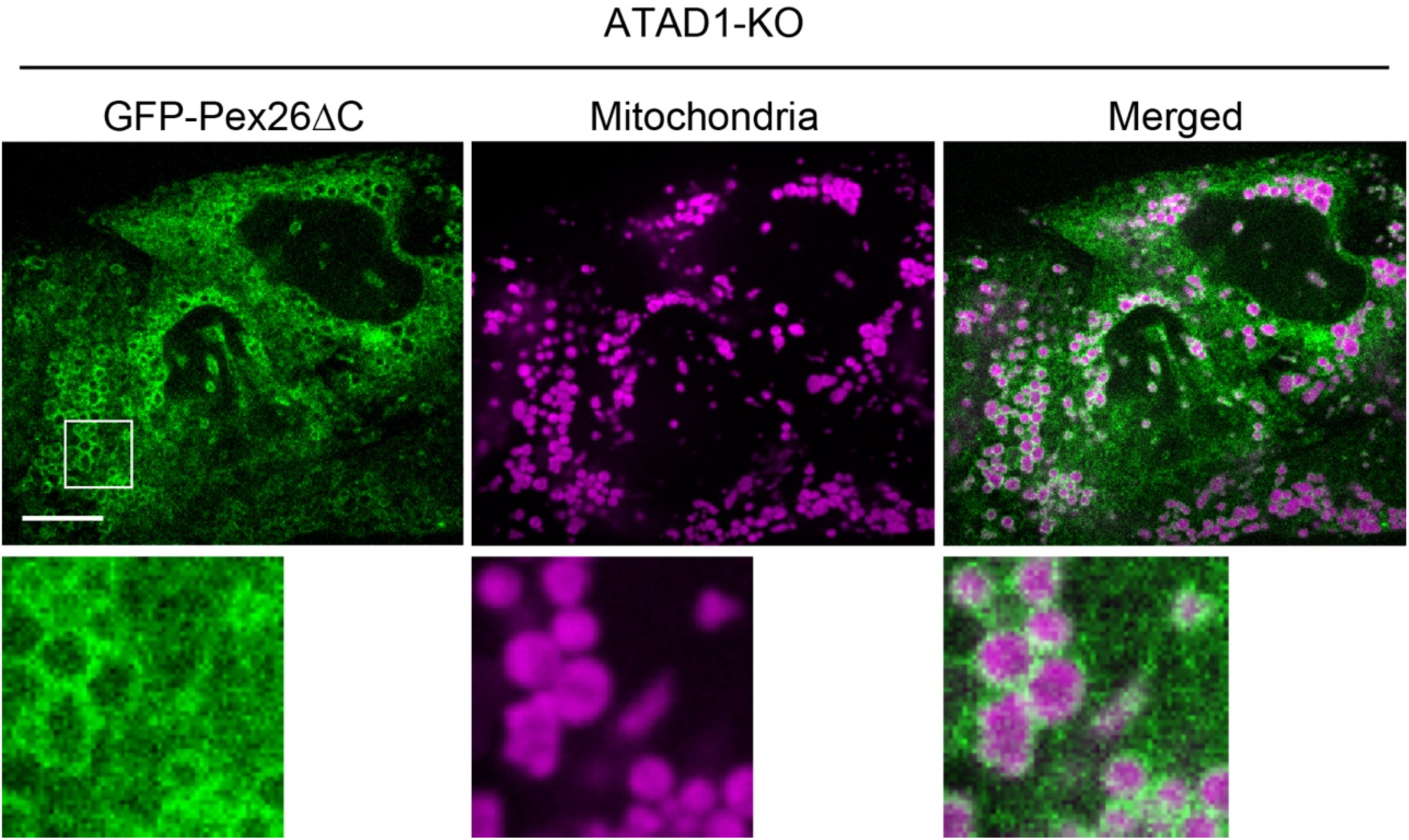
GFP-Pex26ΔC localization in MARCH6-depleted ATAD1-KO cells. ATAD1-KO cells stably expressing GFP-Pex26ΔC were treated with MARCH6 siRNA and analyzed by confocal microscopy. Mitochondria and the ER were visualized with MitoTracker and mCherry-Sec63, respectively.

**Table S1.** Plasmids used in this study.

| Plasmid # | Plasmid Name | Reference |
|---|---|---|
| pMM451 | pcDNA3.1-Tom70N-FLAG-sTurboIDn-GFP1-10 | This study |
| pMM449 | pcDNA3.1-ERj1N-V5-GFP11-sTurboIDc | This study |
| pMM443 | pcDNA3.1-ERj1N-V5-sTurboIDc-GFP11 | This study |
| pMM450 | pcDNA3.1-Tom70N-FLAG-GFP1-10-sTurboIDn | This study |
| pMM441 | pcDNA3.1-Tom70N-FLAG-GFP11-sTurboIDn | This study |
| pMM444 | pcDNA3.1-ERj1N-V5-GFP1-10-sTurboIDc | This study |
| pMM445 | pcDNA3.1-ERj1N-V5-sTurboIDc-GFP1-10 | This study |
| pMM442 | pcDNA3.1-Tom70N-FLAG-sTurboIDn-GFP11 | This study |
| pMM82 | pCDNA3.1-Tom70N-FLAG-GFP1-10 | Kakimoto et al., 2018 |
| pMM86 | pCDNA3.1-ERj1N-V5-GFP1-10 | Tashiro et al., 2020 |
| pMM82 | pCDNA3.1-Tom70N-FLAG-GFP1-10 | Kakimoto et al., 2018 |
| pMM87 | pCDNA3.1-ERj1N-V5-GFP11 | Kakimoto et al., 2018 |
| pMM475 | pcDNA3.1-3XHA TurboID | This study |
| pKO95 | ATAD1KO-CHOP3 | This study |
| pMM713 | pcDNA3.1-Pex3(1-33)-mCherry | This study |
| pMM480 | pcDNA3.1-mCherry-sec61b | This study |
| pMM243 | pcDNA3.1-Su9-tagBFP | This study |

**Table S2.** Antibodies used in this study.

| Antibody |  | Cat. No. | RRID |
|---|---|---|---|
| V5 | MBL | M-215-3 | AB_2811168 |
| FLAG | Sigma-Aldrich | F1804 | AB_262044 |
| GFP | Takara Bio | 632381 | AB_10013427 |
| VAPA | Proteintech | 15275-1-AP | AB_2256991 |
| VAPB | Proteintech | 14477-1-ap | AB_2288297 |
| RMDN3 | Proteintech | 20641-1-ap | AB_10695187 |
| BAP31 | Proteintech | 11200-1-ap | AB_2065142 |
| ATAD1 | Proteintech | 16882-1-AP | AB_2878327 |
| RHOT1 | Sigma-Aldrich | WH0055288M1 | AB_1843347 |
| RHOT2 | Proteintech | 11237-1-AP | AB_2179539 |
| TOMM40 | Proteintech | 18409-1-AP | AB_2303725 |
| Mfn1 | Proteintech | 13798-1-AP | AB_2266318 |
| Mfn2 | Santa cruz | sc-515647 | AB_2811176 |
| Grp75 | Proteintech | 14887-1-AP | AB_2120458 |
| VDAC | abcam | ab15895 | AB_2214787 |
| IP3R3 | BD Bioscience | 610312 | AB_397704 |
| RAB7A | Proteintech | 55469-1-AP | AB_11182831 |
| PDZD8 | LSBio | LS-C345961 |  |
| SYNJ2BP | Proteintech | 15666-1-AP | AB_2201149 |
| actin | Santa Cruz Biotechnology | sc-47778 | AB_626632 |
| TOM20 | BD transduction laboratories | 612278 | AB_399595 |
| Cy5 AffiniPure Goat Anti-Mouse IgG (H+L) | Jackson ImmunoResearch Laboratories | 115-175-166 | AB_2338714 |
| DyLight 800-conjugated goat anti-rabbit IgG (H+L) | Rockland Immunochemicals | 611-145-002-0.5 | AB_11183542 |
| Goat anti-Mouse IgG (H+L) Cross-Adsorbed Secondary Antibody, Alexa Fluor™ 405 | Thermo Fisher Scientific | A48255 | AB_221604 |
| Streptavidin - Cy5 | Thermo Fisher Scientific | 434316 |  |

**Table S3.**
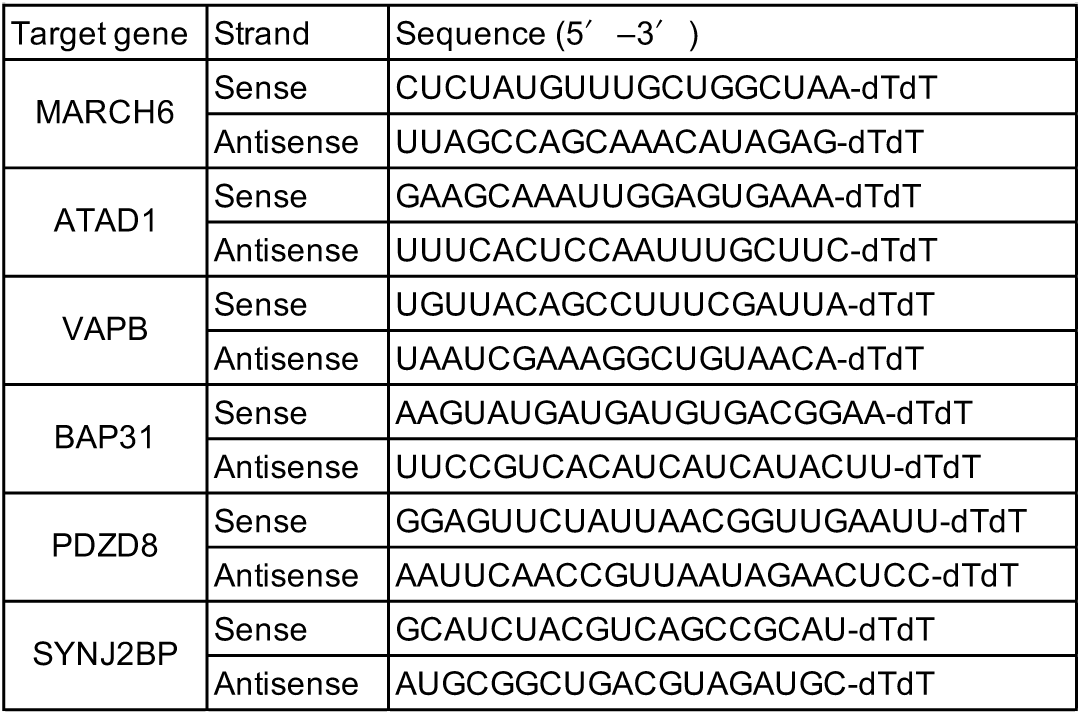
siRNA sequences used in this study.

